# Decoding the role of microbial interspecies interactions on nitrogen fixation

**DOI:** 10.64898/2026.08.11.744197

**Authors:** Claire M. Palmer, Jaron Thompson, Jihyun Hanna Hwang, Will Ranger, Jean-Michel Ané, Ophelia S. Venturelli

## Abstract

Nitrogen fixation performed by rhizosphere bacteria has the potential to improve the sustainability of cereal crop cultivation. Deciphering the role of interspecies interactions on nitrogen fixation is crucial for devising strategies to enhance this process. To unravel the contributions of interspecies interactions, we constructed synthetic microbial communities from the bottom-up that contain diazotrophic bacteria that fix nitrogen and maize rhizosphere bacteria that do not have this capability. Interactions that impacted nitrogenase activity via growth-independent mechanisms were prevalent in the system. Nitrogenase activity increased and eventually saturated as a function of the number of inoculated diazotrophs. Using a tailored machine learning model for microbiome dynamics and explainable artificial intelligence, we deciphered species contributions on nitrogenase activity and diazotroph growth. We identified a community containing *Klebsiella variicola*, *Herbaspirillum seropedicae*, and *Stutzerimonas stutzeri* as a starting point for developing microbial inoculants for cereal crops. Taken together, these results provide insights into the role of interspecies interactions on nitrogenase activity.

## INTRODUCTION

Microbial communities are present in nearly every environment, including soil, the rhizosphere, and the human gut^1,2^. A major goal is to predict and manipulate microbiome functions, which are vital to host health and every ecosystem on Earth^3–5^. One such valuable function is biological nitrogen fixation performed by diazotrophic members of the rhizosphere microbiome^6–8^. Diazotrophs use the nitrogenase enzyme to convert dinitrogen gas into ammonia. The fixed nitrogen can then be released as ammonium and used by the plant host^9–11^. Associative diazotrophs (species that live close to the roots of non-leguminous plants) are of particular interest for cereal crops such as maize, sorghum, and rice, which rely heavily on synthetic nitrogen fertilizers^12–14^. While these fertilizers enable the production of nearly half of the world’s food supply^6^, they have negative environmental consequences^13,14^ and fluctuate in cost, creating a financial burden for growers^15,16^. Microbial inoculants could provide fixed nitrogen to cereal crops in a sustainable manner^17^. However, interspecies interactions within the rhizosphere microbiome represent a key control knob that remains poorly understood.

The regulatory mechanisms controlling nitrogen fixation have been extensively studied in diazotrophs isolated from cereals or soil, leading to the generation of mutants that are insensitive to the presence of nitrogen, allowing for the overproduction and release of ammonia^17–20^. The application of both engineered and wild-type diazotrophic strains has been shown to benefit cereal growth^21–25^. However, further enhancement and discovery of the limits of applying nitrogen-fixing microbial inoculants necessitate a deeper understanding of the ecological and molecular factors that drive inoculant establishment, persistence, and impact on the resident microbiome^26,27^. While improving inoculant growth could enhance the desired functions, the quantitative relationship between diazotroph growth and nitrogenase activity has not yet been resolved.

Communities that provide fixed nitrogen to cereal crops naturally exist. For example, maize and sorghum with a unique aerial root mucilage-producing phenotype can obtain around half of their nitrogen from associated diazotrophs^28,29^. Maize xylem sap also hosts nitrogen-fixing communities^30^. While these communities can benefit the plant, we have a limited understanding of how interactions within these communities impact nitrogen fixation. A previous study demonstrated that a four-member synthetic community containing two diazotrophs and two non-diazotrophs had higher nitrogenase activity than the two diazotrophs alone^30^. This implies that in this case interspecies interactions enhanced nitrogenase activity. However, the distributions of beneficial versus antagonistic interactions within rhizosphere microbial communities are unknown.

Synthetic microbial communities enable precise control of the initial abundance of each species^3,4,31^. Computational models informed by high-throughput experimental measurements of synthetic communities have been used to predict community dynamics, functions, and to uncover interspecies interaction networks^32–34^. Machine learning models can capture species growth and other variables of interest (e.g. metabolites) and can outperform mechanistic models such as the generalized Lotka-Volterra and consumer-resource models at prediction tasks^35–37^. Bayesian optimization is a principled method to enable efficient exploration of high-dimensional design spaces^36^. This approach has been applied to design and optimize microbial communities with desired functions by sampling a subset of highly informative conditions to fill knowledge gaps in the model and optimize desired objectives^38^. Explainable artificial intelligence (AI) methods, such as Shapley additive explanations (SHAP), can be used to decipher microbial interactions using a trained and validated model^38,39^.

To provide insights into the prevalence of interspecies interactions that impact nitrogen-fixing activity, we performed detailed and quantitative characterization of the growth and nitrogen-fixing activity of a synthetic community comprising five associative diazotrophs^40–42,20,43^ and seven non-diazotrophs isolated from maize roots^44^. Using a machine learning model tailored for microbiomes informed by experimental data^36^, we revealed key insights into the impact of interspecies interactions on nitrogenase activity in this community. Diazotroph growth was not the sole driver of nitrogenase activity, demonstrating that species can impact nitrogenase activity through growth-uncoupled mechanisms. While most interspecies interactions negatively impacted nitrogenase activity, we identified sparse interactions that positively influence nitrogenase activity and determined that a particular beneficial interaction can be replicated by conditioned media experiments. Nitrogenase activity increased and saturated as a function of the number of inoculated diazotrophs. We identified key contributors to community nitrogenase activity using explainable artificial intelligence. Finally, we used this information to select a high-activity community for the exploration of the minimal number of species necessary to achieve high nitrogenase activity. Taken together, these results provide a framework for the rational design of microbiome-based strategies to enhance nitrogen fixation in the cereal rhizosphere and offer insights into the relationships between interspecies interactions and nitrogen fixation more broadly.

## MATERIALS AND METHODS

### Strain, media, and culturing conditions

The strains used in this work were obtained from the sources listed in Table S1. Strains were routinely cultured aerobically at 30℃ in TY medium except Ab and Av, which were cultured in TY medium with the addition of 20 g/L sucrose (TY+sucrose). Permanent stocks of each strain were stored in 25% glycerol at -80℃. Batches of single-use glycerol stocks were produced for each strain by first isolating single colonies from the permanent stock on solid TY or TY+sucrose media and then inoculating liquid culture with a single colony and growing it to the late exponential phase. Cultures were then mixed in equal volume of 50% glycerol, 400 μl of the mixture was aliquoted into Matrix Tubes (ThermoFisher) and stored at -80℃. The identity of the organism was verified by 16S rRNA gene sequencing. All experiments were performed in a chemically defined medium, CMP-ARE, the composition of which is provided in Table S2. This medium was developed based on a previously studied maize artificial root exudate medium^45^ and Burk’s medium for nitrogen-fixing species^46^. Microaerobic culturing was performed in a hypoxic chamber (Coy Labs) with an atmosphere of 3±0.2% O_2_, 0.2±0.1% CO_2_, and a balance of N_2_.

### Community culturing experiments

For each experiment, we utilized a two-stage preculturing setup. First, 5 mL of TY or TY+sucrose medium was inoculated with 100 μL of a thawed single-use glycerol stock and incubated aerobically with shaking at 30°C. Next, the volume of preculture listed in Table S3 was inoculated into 5 mL of CMP-ARE medium (multiple tubes were inoculated if a larger volume was needed) and incubated at 30℃ with shaking under microaerobic conditions, except for Av, which was grown aerobically. The timing of each preculturing step for each species is listed in Table S3. Following this second growth step, the preculture of each species was diluted to an optical density at 600 nm (OD_600_) of 0.05 using a Tecan F200 Plate Reader (200 μl in a 96-well microplate) in CMP-ARE medium. Community combinations were arrayed in 2.2 mL, 96-deepwell plates by pipetting equal volumes of each species’ diluted preculture into the appropriate wells using a Tecan Fluent Liquid Handling Robot inside a hypoxic chamber. Assembled communities were diluted in triplicate into 2.2 mL, 96-deepwell plates to a final volume of 1 mL and an OD_600_ of 0.01. Each plate was covered with a semi-permeable membrane (Breathe-Easy®) and incubated at 30°C in the dark with shaking in a hypoxic chamber. After 16, 24, or 48 hours, plates were removed from the incubator. The OD_600_ of 200 μl of each culture was measured in a 96-well microplate in a Tecan F200 Plate Reader. Samples were collected by transferring 200 μl each to a 1 mL round-bottom, 96-deepwell plate, centrifuging at 2400 × g for 15 minutes. 20 μL of each supernatant was used to quantify the pH using a phenol red assay as described in Clark 2021^34^. Cell pellets were stored at -80°C for subsequent genomic DNA extraction.

### Acetylene reduction assay

For acetylene reduction assays, assembled communities were prepared in a hypoxic chamber as previously described and diluted into 10 mL vials (Supelco) to a final volume of 1 mL and an OD_600_ of 0.01. Vials were sealed with crimp caps (MicroSolv #95025-02-1S), removed from the hypoxic chamber, and incubated at 30°C, in the dark, with shaking. After 44 hours, 1 mL of air in the headspace was removed from each vial using a syringe fitted with a gauge 26 needle (BD), 1 mL of acetylene (99% v/v) was injected, and the vials were returned to incubation under the same conditions. After an additional 4 hours, 1 mL gas samples were removed from each vial using a syringe, inserted into a large rubber stopper, and stored at room temperature for future analysis. The vials were then opened, and the culture was transferred to a 96-deepwell plate. OD_600_ measurements were taken and samples collected as previously described. Gas samples were later injected into fresh, capped 10 mL vials (Supelco, MicroSolv cap) after removing 1 mL of headspace air, and analyzed using a GC-2010 instrument (Shimadzu) equipped with a 60 m Rt-Alumina BOND/KCL column (Restek). Ethylene in each vial was quantified using a standard curve, with the limit of detection being 0.3 nanomoles. The nanomoles of ethylene in each original sample vial was calculated by multiplying the measured ethylene by 10 to account for the dilution. Ethylene production per hour was calculated by dividing the ethylene in the sample by the incubation time with acetylene. Relative nitrogenase activity was calculated by dividing the ethylene production per hour by the mean ethylene production per hour of the entire community on the same experimental day.

### Genomic DNA extraction, library preparation, Illumina sequencing, and analysis

DNA extraction, library preparation, sequencing, and analysis were performed as described in Hromada 2021 and Clark 2021^33,34^.

### The Microbiome Recurrent Neural Network (MiRNN)

To model the dynamics of species abundance, relative nitrogenase activity, and pH, we used the Microbiome Recurrent Neural Network (MiRNN), a machine learning model tailored to predict microbial community dynamics that eliminates the possibility of predicting physically unrealistic species abundances. The MiRNN can be trained to predict species abundance, relative nitrogenase activity, and pH over time in response to community composition. Once trained, the MiRNN can be used to select experimental conditions predicted to maximize relative nitrogenase activity. The MiRNN is described in full in Thompson 2023^36^.

### Parameter estimation

To estimate model parameters, θ, we use an approximate Bayesian inference method to infer a parameter distribution by conditioning on experimentally observed data. We define a set of n experimental conditions as *Q* = {q_1_, …, q_n_} and a set of corresponding measurements of species, pH, and relative nitrogenase activity as *Ɗ*(*Q*) = {y(q_1_), …, y(q_n_)}. Measurements of relative nitrogenase activity were preprocessed such that any sample containing diazotrophic species that did not produce ethylene above the limit of detection was treated as a missing value rather than a true zero. We use the Laplace approximation to compute the mean and covariance of a Gaussian distribution that approximates the true parameter posterior. A prior parameter distribution centered at zero is used to promote model simplicity, denoted as p(θ) = *Ɲ*(0, α^−1^Ⅱ_n_θ__). The precision of the prior, α, is a hyperparameter that determines how much the parameters are penalized for deviating from zero.

Because measurements of species, pH, and relative nitrogenase activity are affected by many potential sources of noise, we assume the distribution of measurement noise is Gaussian. We observed that the standard deviation across replicate measurements of both relative nitrogenase activity and species abundances was significantly correlated with the replicate mean (**Figure S1**). We therefore model the distribution of species and relative nitrogenase activity conditioned on a particular experimental condition q as a Gaussian with a mean predicted by the MiRNN and with a standard deviation that scales in proportion to the predicted mean,

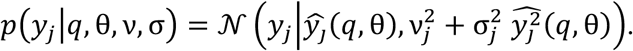

where j = 1, …, n_s_ + n_m_ indexes a particular species or metabolite. Given this model for the distribution of species and metabolites, the likelihood of observing a particular set of measured values *Ɗ*(*Q*) is

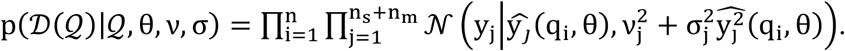

Bayes theorem defines the posterior parameter distribution as proportional to the likelihood of a dataset multiplied by the prior distribution over parameters,

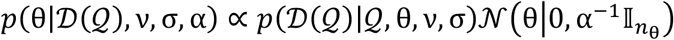

Maximizing the log of the parameter posterior distribution with respect to θ gives the Maximum A Posteriori (MAP) estimate,

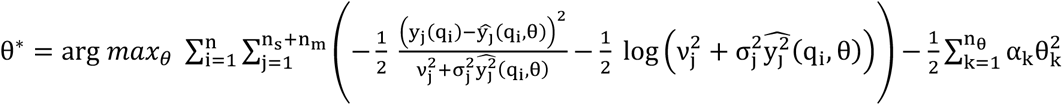

The Laplace method approximates the posterior parameter distribution as a Gaussian with a mean of θ^∗^ and covariance as the inverse of the matrix of second derivatives of the negative log posterior,

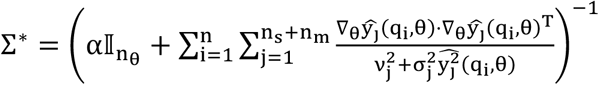

With the posterior parameter distribution approximated as a Gaussian, p(θ|*Ɗ*(*Q*), ν, σ, α) = *Ɲ*(θ|θ^∗^, Σ^∗^), the posterior predictive distribution is determined by marginalizing with respect to the posterior,

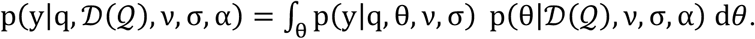

We use the Expectation-Maximization (EM) algorithm to optimize the precision of the parameter prior, α^36,47^. Optimization of measurement precision parameters for each predicted variable, ν_j_ and σ_j_, is performed by keeping θ fixed to the MAP estimate and maximizing the log likelihood function,

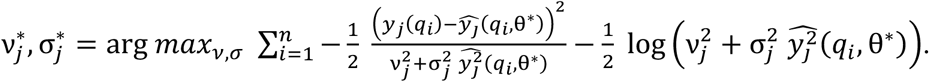

### Bayesian optimization

Once the MiRNN is trained on previously collected data, we can use the model to predict which species combinations will result in increased relative nitrogenase activity. Thompson sampling is a Bayesian optimization algorithm that exploits and explores an objective function by first sampling candidate parameter values from the posterior parameter distribution, θ^(k)^ ∼ *Ɲ*(θ^∗^, Σ^∗^), and then uses the resulting model to identify the condition that optimizes a predicted objective^48^. For each parameter sample, k = 1, …, K, an experimental condition is selected by maximizing predicted relative nitrogenase activity,

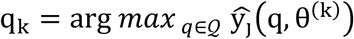

where the index j in this case corresponds to relative nitrogenase activity. The process of sampling and selecting a condition is repeated until the desired number of conditions is selected.

### Statistical analyses

All statistical analyses were performed using Python version 3.12.2. Spearman correlations, Pearson correlations, and Dunnett’s tests were performed using the stats module of the Python package SciPy. Levene tests, ANOVA, Welch ANOVA, and Games-Howell tests were performed using the Python package Pingouin. Root mean squared error was calculated using Python package scikit-learn. Plots were made using the Python packages Seaborn and Matplotlib.

## RESULTS

### A synthetic community to study the role of interspecies interactions on nitrogen fixation

We constructed synthetic microbial communities from the bottom-up, containing associative diazotrophic species and non-fixing bacteria isolated from the cereal rhizosphere (**Figure 1A**, **Table S1**). In particular, we included a seven-member bacterial community that has been shown to stably and reproducibly assemble on maize roots^44^ and has been used as a model of the maize rhizosphere microbiome^49–52^. This community comprises Pseudomonadota (5 species), Bacteroidota (1 species), and Actinomycetota (1 species), which are prevalent and abundant organisms in the rhizosphere and constitute the Pseudomonadota-dominated taxonomic profile in this environment^2,44^. To enable nitrogen fixation, we introduced three well-studied associative diazotrophs^40–42^ and two recently isolated nitrogen-fixing strains^20,43^ to establish a 12-member nitrogen-fixing community. Evaluation of nitrogen fixation is typically performed in diazotrophic media containing a single carbon source and no nitrogen source^46^. However, the cereal rhizosphere can include a mixture of carbon sources, including sugars, organic acids, and amino acids^45^. To mirror this nutrient complexity, we combined an artificial maize root exudate^45^ and a diazotrophic growth medium^46^ to generate a new medium that supported the growth of all individual species and nitrogenase activity in the 12-member community (**Figure S2**).

**Figure 1.**
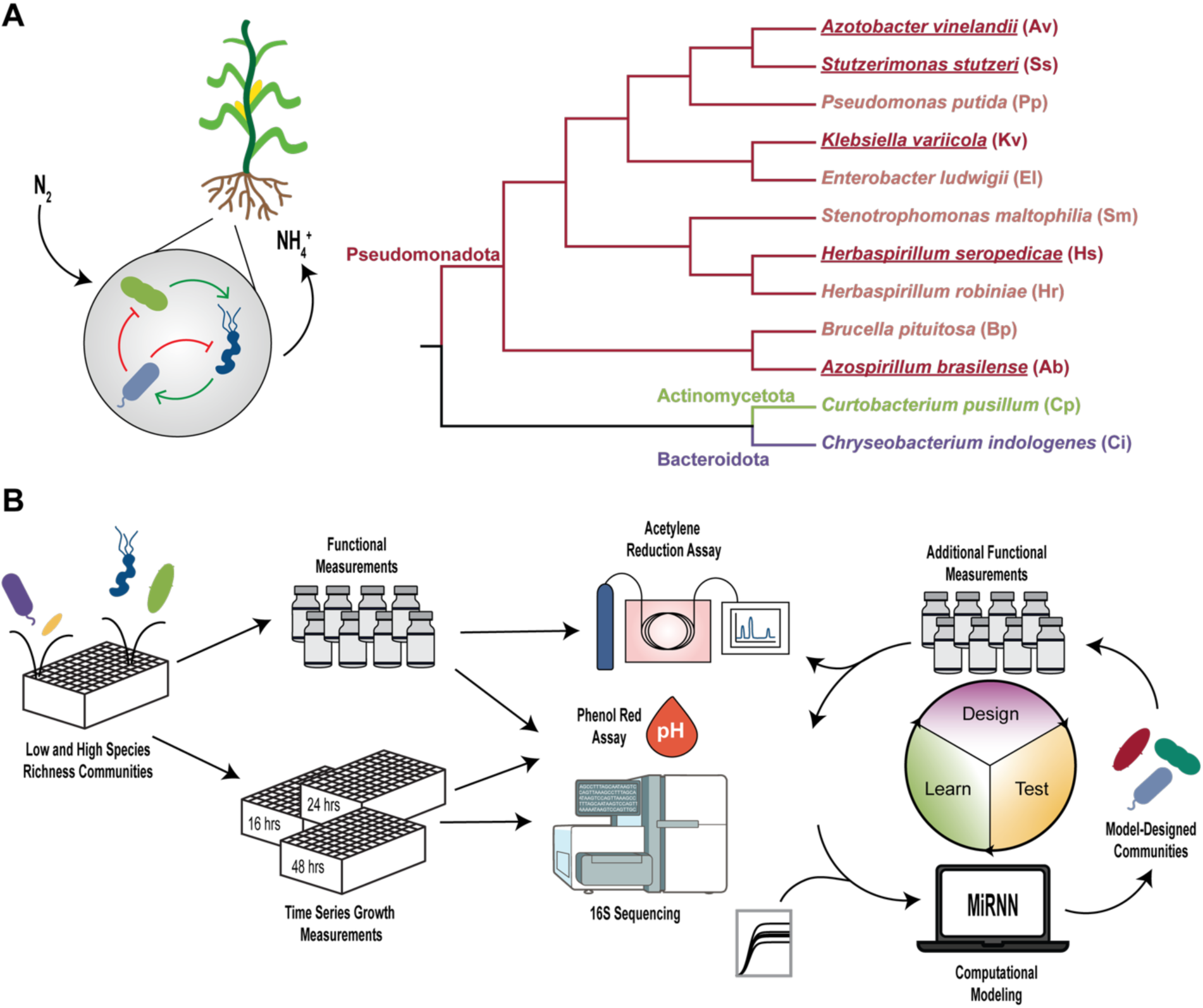
Investigation of interspecies interactions on nitrogen fixation using a synthetic rhizosphere community. A) Phylogenetic tree of a 12-member synthetic rhizosphere community based on a concatenated alignment of 392 marker proteins, using PhyloPhlAn version 3.0^53^. Branch color indicates phylum, and diazotrophs are underlined. B) Schematic representing the experimental workflow. For initial experiments, synthetic communities were assembled under hypoxic conditions and inoculated into both vials for acetylene reduction assays and deep-well plates for time-series measurements. The absolute abundance of each species was determined by measuring cell density at 600 nm and community composition using multiplexed 16S rRNA gene sequencing. The pH of each sample was determined through phenol red assays. The functional dataset was further expanded using a design-test-learn cycle. Functional and absolute abundance data were used to train a Microbiome Recurrent Neural Network (MiRNN). The resulting model was then used to design communities for further experiments.

Sampling the extremes of the community design space using low and high species richness subcommunities has been shown to be an informative starting point for investigating the impact of interspecies interactions on community functions^32^. Therefore, we characterized all possible one-, two-, and 11-species subcommunities of our 12-member community, as well as five rationally designed communities of interest, such as the seven-member maize rhizosphere community (**Figure 1B**, left). We assessed nitrogenase activity by measuring the nitrogenase-catalyzed reduction of acetylene to ethylene, in conjunction with community composition via 16S rRNA gene sequencing and total community biomass (OD_600_). We measured endpoint extracellular pH because variation in environmental pH is a key factor shaping the rhizosphere microbiome^2^. We assessed community composition and extracellular pH at 16, 24, and 48 hours (**Figure 1B**, left) and nitrogenase activity at a single timepoint due to the low throughput of acetylene reduction assays. Overall, this dataset consisted of functional and time-resolved measurements of 96 unique monocultures and community combinations.

The full 12-member community served as a reference control for all experiments, controlling for growth and metabolic variability in the pre-cultures. Therefore, we normalized nitrogenase activity for each community to the full community’s nitrogenase activity in the same experiment. We refer to this normalized value as relative nitrogenase activity. This dataset was used to inform the selection of 30 medium-richness communities to expand the experimental characterization of this sparsely sampled richness region of the design space (**Figure 1B**, right).

We leveraged Bayesian optimization to maximize the information value in additional experiments since we were unable to exhaustively explore the community design space (4,095 possible species combinations). To combine insights from our time-resolved dataset and our expanded functional dataset, we used a flexible machine learning model referred to as the Microbiome Recurrent Neural Network (MiRNN)^36^ (**Figure S1**). For the next experimental cycle, 30 communities were selected to balance exploration and exploitation (maximize relative nitrogenase activity). The resulting dataset was used to train an updated MiRNN informed by 156 unique communities, with time-resolved measurements from 96 communities and nitrogenase activity measurements from 149 communities. Using 20-fold cross-validation, the MiRNN model displayed a Pearson correlation of 0.79 (p-value = 1.81×10^-72^) and a Spearman correlation of 0.89 (p-value = 3.17 ×10^-116^) for relative nitrogenase activity (**Figure S3M**, **Table S4**). To further explore the community design space, the MiRNN trained on all data was used to predict the relative nitrogenase activity for all possible community combinations, yielding an *in silico* predictions for the full design space of 4,095 communities.

### Nitrogenase activity is not driven solely by diazotroph growth

Interspecies interactions that modulate diazotroph growth have the potential to impact nitrogenase activity. Across our experimental dataset, diazotroph abundance was positively correlated with relative nitrogenase activity (Spearman r = 0.72, p-value = 1.59×10^-25^). However, certain communities displayed a high endpoint diazotroph abundance and low relative nitrogenase activity (low species richness communities in the bottom right quadrant, **Figure 2A**). By contrast, other communities exhibited high nitrogenase activity and moderate diazotroph abundance (medium to high species richness communities in the top-left quadrant). To go beyond our experimental measurements, we used the MiRNN model to predict nitrogenase activity and diazotroph abundance in all possible communities (**Figure 2B**). In communities containing the diazotroph *Klebsiella variicola* (Kv), predicted diazotroph abundance was positively correlated with predicted relative nitrogenase activity (Spearman r = 0.48, p-value = 1.29×10^-173^). Despite this, some communities predicted to have high diazotroph abundance had low predicted nitrogenase activity (bottom-right quadrant). Other communities predicted to have high nitrogenase activity only had moderate predicted diazotroph abundance (top-left quadrant). Further, in communities lacking Kv there was no significant correlation (Spearman r = 0.04, p-value = 0.06). Taken together, these results suggest that diazotroph growth is not the sole driver of community nitrogenase activity and interspecies interactions may impact nitrogenase activity through growth-uncoupled mechanisms. In selected pairwise communities containing a single diazotroph, only one species can directly contribute to nitrogenase activity, revealing interspecies interactions that impact nitrogenase activity of each diazotroph. Several cocultures exhibited a significant decrease in the relative nitrogenase activity compared to the monoculture, demonstrating antagonistic interactions that impact nitrogenase activity (**Figure 2C**). In particular, the diazotroph, *Stutzerimonas stutzeri* (Ss), had significantly lower relative nitrogenase activity in coculture with six of seven non-diazotrophs. In coculture with the non-diazotroph *Enterobacter ludwigii* (El), the abundance of Ss significantly decreased compared to monoculture, suggesting a growth-mediated mechanism of inhibition (**Figure 2D**). However, in other pairwise communities containing Ss with significantly lower relative nitrogenase activity, the absolute abundance of Ss did not change significantly compared to the monoculture. Similarly, the nitrogenase activity of Kv substantially decreased in coculture with the non-diazotroph, *Brucella pituitosa* (Bp) whereas Kv’s abundance did not change significantly. Kv acidified the growth medium across different community contexts (**Figure S4**). However, the Bp and Kv pairwise community displayed a significant increase in endpoint pH compared to the Kv monoculture (**Figure S5E**). Therefore, negative impacts of interspecies interactions on nitrogenase activity were often growth-independent with six of seven negative interactions on nitrogenase activity having no effect on growth.

**Figure 2.**
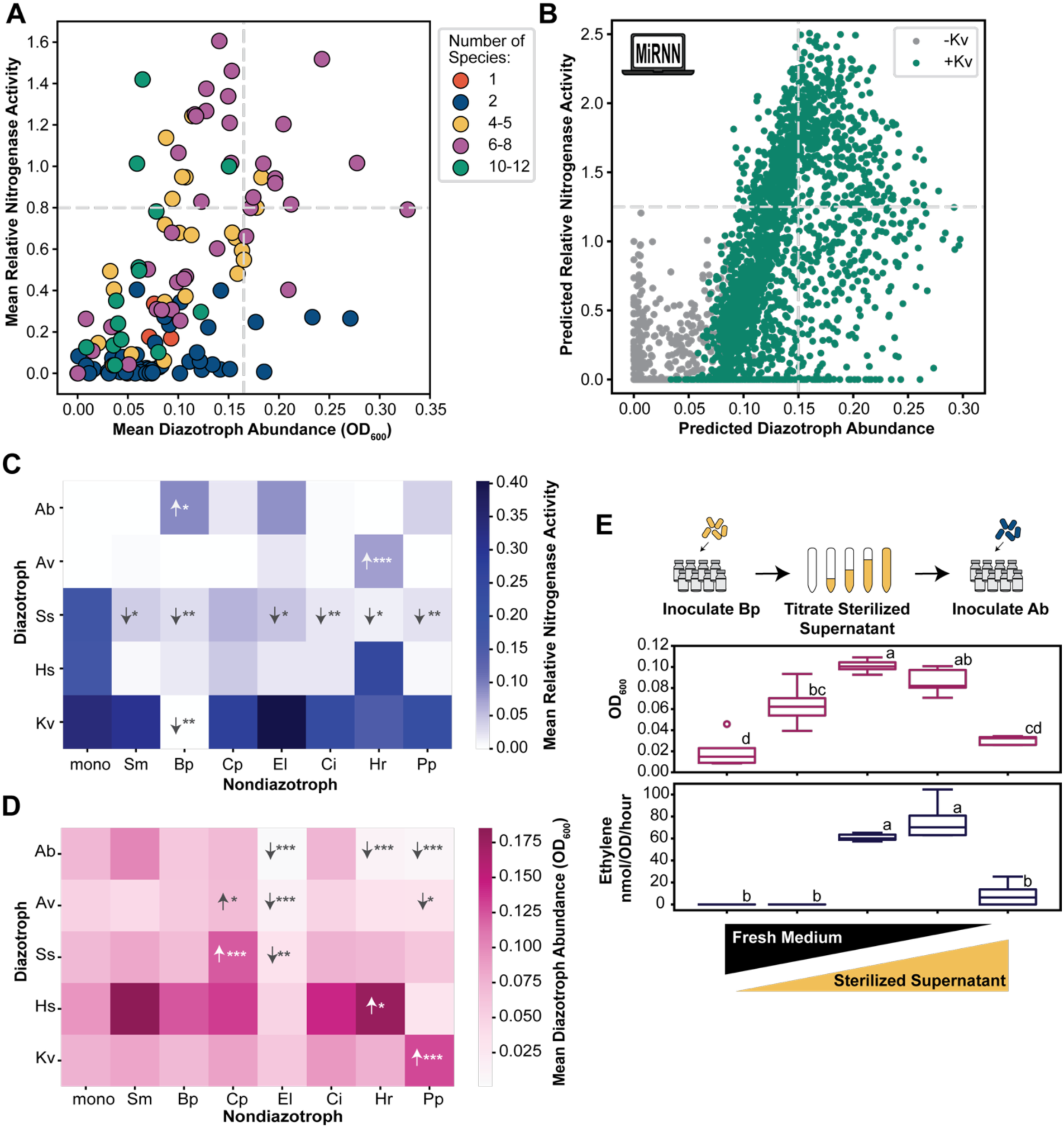
Interspecies interactions can modulate nitrogenase activity independent of diazotroph growth. A) Relative nitrogenase activity versus diazotroph abundance for each community in the experimental dataset. Diazotroph abundance is calculated by summing the absolute abundance of each diazotrophic species. Each data point represents a unique community, with color representing the total number of species in the community. Dashed lines indicate the halfway point of each axis. B) Predicted relative nitrogenase activity versus predicted diazotroph abundance for each community in the simulated dataset. Each data point represents a unique community, with color representing the presence or absence of diazotroph, *Klebsiella variicola*. Dashed lines indicate the halfway point of each axis. Heatmaps comparing the C) relative nitrogenase activity and D) diazotroph abundance of each diazotroph in monoculture to pairwise cocultures with each non-diazotrophic species. For each diazotroph, pairwise cocultures were compared with the monoculture control using Dunnett’s test when variances were equal (as determined by the Levene test), and the Games-Howell test when variances were unequal. * p < 0.05, ** p < 0.01, *** p < 0.005 E) Growth and ethylene production of Ab monoculture in (from left to right), 100% fresh medium, 75% fresh medium and 25% Bp sterilized supernatant, 50% fresh medium and 50% Bp sterilized supernatant, 25% fresh medium and 75% Bp sterilized supernatant, and 100% Bp sterilized supernatant. Boxplot hinges indicate the first and third quartiles. Biological replicates outside of 1.5 times the interquartile range are indicated by circles. Letters groupings statistically distinct groups (p < 0.05) as determined by using Tukey test when variances were equal (as determined by the Levene test), and the Games-Howell test when variances were unequal (n=5). Full statistics are listed in the Extended Statistics section.

Two pairwise communities displayed synergistic interactions that enhanced nitrogenase activity. The diazotroph *Azospirillum brasilense* (Ab) displayed significant enhancement in relative nitrogenase activity in coculture with the non-diazotroph, Bp (**Figure 2C**). Similarly, the presence of *Herbaspirillum robiniae* (Hr) lead to a significant increase in the relative nitrogenase activity of diazotroph, *Azotobacter vinelandii* (Av) (**Figure 2C)**. In both cases, diazotroph abundance did not significantly change (**Figure 2D**). The Ab and Bp pair exhibited a significant reduction in endpoint pH compared to the Ab monoculture (**Figure S5A**). Notably, Bp had the opposite impact on nitrogenase activity and endpoint pH in coculture with Ab as it did with Kv, demonstrating that each diazotroph displays unique interspecies interactions with non-diazotrophs. Finally, while several pairwise communities showed a significant increase in diazotroph growth, none of these enhancements were associated with significant changes in relative nitrogenase activity. Therefore, both synergistic interactions in pairwise communities that enhanced nitrogenase activity arose from growth-independent mechanisms.

To further explore potential mechanisms of interaction of Bp on the diazotrophs Ab and Kv, we performed conditioned media experiments (**Figure 2E and Figure S6**). Intermediate concentrations of the sterilized supernatant of Bp significantly increased the nitrogenase activity of Ab in monoculture compared to the fresh medium control (**Figure 2E**). These results suggest that the positive impact of Bp on the nitrogenase activity of Ab observed in pairwise coculture (**Figure 2C**) is modulated through the extracellular environment. In addition, the sterilized supernatant of Bp also led to a significant increase in the growth of Ab (**Figure 2E**), inconsistent with the trends in the pairwise experiment (**Figure 2D**). By contrast, the sterilized supernatant of Bp did not significantly modify the nitrogenase activity of Kv (**Figure S6A**). The modifications to the extracellular environment by Bp benefited the growth and nitrogenase activity of Ab, however the pairwise experiment demonstrates that this improvement in growth is not required to observe an improvement in nitrogenase activity.

### Nitrogenase activity increases with the number of diazotrophs in the community

In our experimental dataset, relative nitrogenase activity increased and then saturated with the number of inoculated diazotrophs (**Figure 3A**). Saturation of nitrogenase activity can arise from competition for limiting nutrients. While communities containing two diazotrophs displayed significantly higher nitrogenase activity than those containing one diazotroph (**Figure 3A**), this difference was not significant among communities composed solely of diazotrophs (**Figure S7A**). These observations underscore the important role of non-diazotrophs in promoting community nitrogenase activity. Further, the predictions of nitrogenase activity based on the MiRNN model increased with the number of included diazotrophs (**Figure 3B**). However, in contrast to the experimental data, nitrogenase activity did not saturate in the model predictions. This implies that the model does not fully capture potential nutrient limitation or other mechanisms (e.g., the release of metabolic waste products) driving the saturation trend, due to limitations in the number of communities used to inform the model.

**Figure 3.**
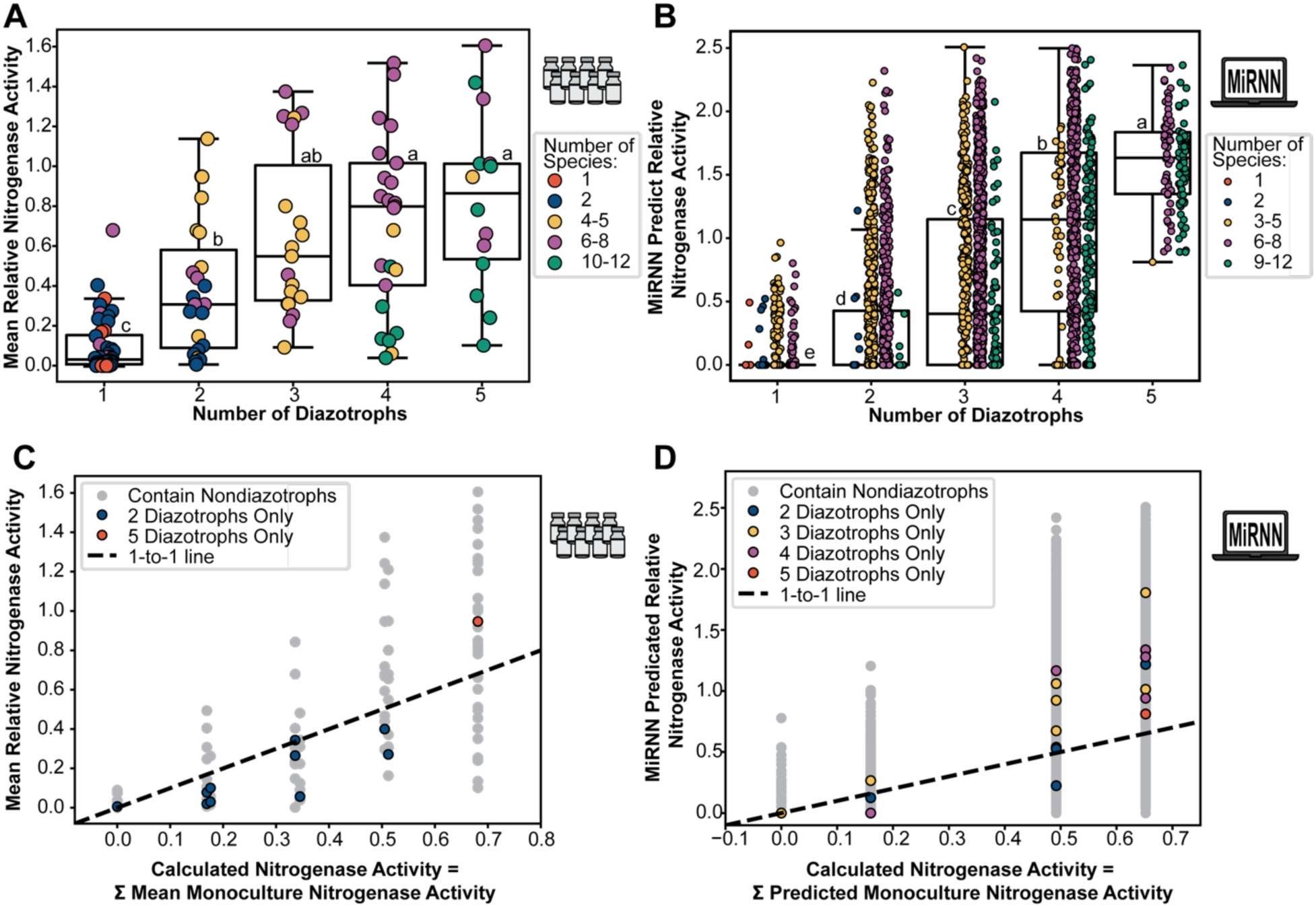
Communities containing multiple diazotrophic species have improved relative nitrogenase activity. Relative nitrogenase activity of each diazotroph-containing community in the A) experimental dataset or B) simulated dataset grouped by the number of diazotrophic species present in the community. Each data point represents a unique community, with color representing the total number of species in the community. Boxplot hinges indicate the first and third quartiles. Letters indicate groupings exhibiting statistically significant relative nitrogenase activity (p < 0.05) as determined by the pairwise Games-Howell test following the Levene test for homoscedasticity and the Welch ANOVA. Complete statistics are listed in the Extended Statistics section. Relative nitrogenase activity of each diazotroph-containing community in the C) experimental dataset or D) simulated dataset, excluding the monocultures, compared to the sum of monoculture nitrogenase activity of each diazotroph in the community. The dashed line represents a one-to-one relationship. Each data point represents a unique community, with color indicating whether the community contains non-diazotrophic species and, for diazotroph-only communities, the number of species.

To explore the deviation from an additive model, we considered a null model in which interspecies interactions are absent. In this case, community nitrogenase activity is the sum of the monoculture nitrogenase activity. We compared the null model with the experimentally measured relative nitrogenase activity in our experimental dataset (**Figure 3C**). While measured nitrogenase activity was correlated with the null model (Pearson r = 0.746, p-value = 1.52×10^-22^), the activity of 31% of communities exceeded the null model (**Figure S7B**). Consistent with our experimental data, 40% of communities were predicted by the MiRNN model to display nitrogenase activity higher than the null model (**Figure 3D**). This implies that synergistic interactions can benefit nitrogenase activity in a range of communities, beyond pairwise cocultures. The null model does not account for resource limitations and thus non-additive synergies may be more substantial when accounting for this major mechanism of interspecies interaction. Diazotroph-only communities with nitrogenase activity exceeding the null model generally included three or more species (**Figures 3C and 3D**). In sum, community nitrogenase activity can be enhanced not only by increasing the number of diazotrophs but also by positive interspecies interactions.

### Modeling insights into nitrogen-fixing community design

To quantitatively capture interactions in the system, we utilized the trained MiRNN model and SHAP^39^ to estimate the effect of each inoculated species on the predicted relative nitrogenase activity in each measured community context (**Figure 4A)**. The diazotroph Kv had the highest median and maximum SHAP values of any species, consistent with its strong contribution to nitrogenase activity across community contexts. The diazotroph, *Herbaspirillum seropedicae* (Hs) also positively contributed to nitrogenase activity across all communities. Av also had a positive impact on nitrogenase activity across many communities but displayed a negative contribution in certain communities. By contrast, the SHAP distributions of diazotrophs Ab and Ss exhibited negative median values, consistent with their low contribution to nitrogenase activity across communities in our experiments.

**Figure 4.**
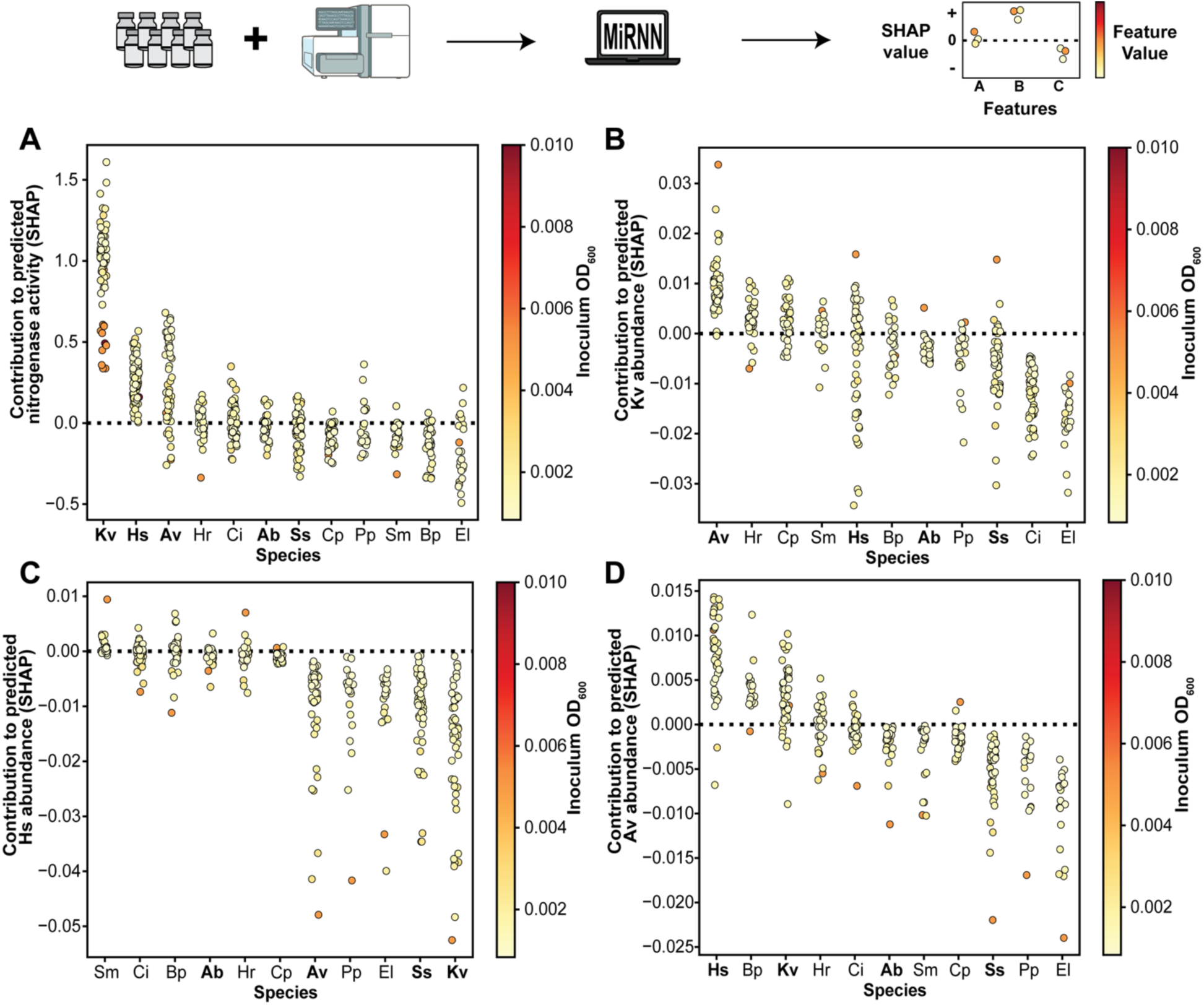
Modelling insights into nitrogen-fixing community design. SHAP values (Shapley additive explanation) of species contribution to model-predicted A) nitrogenase activity, B) Kv abundance, C) Hs abundance and D) Av abundance for each community in the experimental dataset. SHAP values are a game-theory-based approach to explaining each model input’s contribution to a given model output. Species order determined by median SHAP value. Within a species distribution, each point represents a unique combination of species inputs. Color indicates initial inoculum OD_600_ of the species. The dashed line indicates a SHAP value of zero. Diazotrophs are bolded.

Among the non-diazotrophs, most species exhibited negative median SHAP contributions on relative nitrogenase activity, except for *Chryseobacterium indologenes* (Ci) and Hr (**Figure 4A)**. While the SHAP distribution for *Pseudomonas putida* (Pp) showed a negative median value, Pp displayed the highest maximum SHAP contribution of any non-diazotrophic species. The SHAP distributions of all non-diazotrophs exhibited a mixture of positive and negative values, consistent with their context-dependent effects on different diazotrophs (**Figure 4A**). To investigate growth-mediated effects, SHAP was used to determine the effect of each species on the predicted abundance of Kv (**Figure 4B**), Hs (**Figure 4C)**, and Av (**Figure 4D**). While most the species had negative impacts on growth, Av had a positive impact on the growth of Kv. Additionally, Hs and Kv positively impacted the growth of Av, suggesting there are beneficial growth interactions coupling these species. The variation in the distribution of nitrogenase activity for communities containing each diazotroph can provide insight into the robustness of their functional activities across environmental contexts (**Figure S8**). The nitrogenase activity of communities containing Kv varied less than communities containing other diazotrophs in both the experimental and simulated datasets. These results suggest that the nitrogenase activity of Kv was more robust to environmental variation than the other diazotrophs.

### Designing a minimal community with high nitrogenase activity

To provide insight into the species presence/absence that may maximize nitrogenase activity, we grouped communities in our experimental dataset based on their composition in a hierarchical manner. We first selected Kv and then Hs based on the ranking of their median SHAP values. In addition, Bp displayed large negative SHAP values in most communities (**Figure 4A**), and the Bp-Kv pair displayed low nitrogenase activity (**Figure 2A**), suggesting that this species should be excluded from communities to maximize nitrogenase activity. We therefore chose to focus on communities containing Kv and Hs but lacking Bp (+Kv, +Hs, -Bp) as group of interest. Communities in this group had significantly higher relative nitrogenase activity than communities lacking Kv (-Kv) or Hs (+Kv, -Hs) (**Figure 5A**). Among communities containing both Kv and Hs, nitrogenase activity was higher in the absence than presence of Bp, but the difference was not statistically significant (p-value = 0.06). This result indicates that, although some communities can overcome the detrimental impact of Bp’s presence, Bp has an overall negative impact on nitrogenase activity.

**Figure 5.**
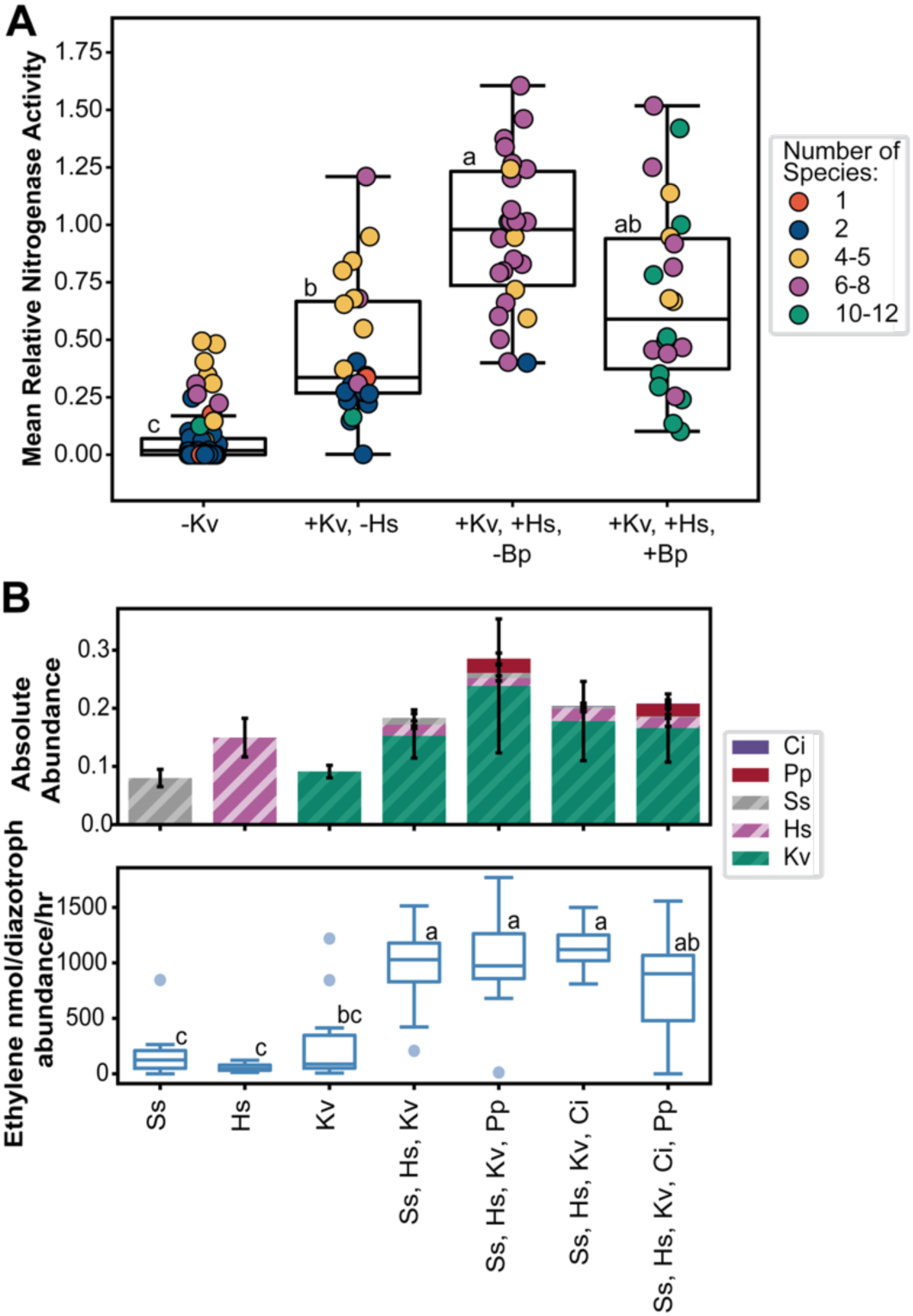
Designing a minimal community with high nitrogenase activity. A) Mean relative nitrogenase activity of all communities, grouped by the presence or absence of key species. Data points represent the mean of biological replicates for each community. Boxplot hinges indicate the first and third quartiles. Letters indicate groupings exhibiting statistically significant relative nitrogenase activity (p < 0.05) as determined by the pairwise Games-Howell test following the Levene test for homoscedasticity and the Welch ANOVA. Complete statistics are listed in the Extended Statistics section. B) The top panel shows community composition after 48 hours. Color represents species, with diazotrophs indicated by hashes. Error bars represent one standard deviation from the mean of biological replicates. Absolute abundance is calculated as the product of relative abundance and cell density at 600 nm. The bottom panel shows community nitrogenase activity, normalized by the sum of the absolute abundance of all diazotrophic species and the number of hours incubated with acetylene. Boxplot hinges indicate the first and third quartiles. Biological replicates with ethylene production outside of 1.5 times the interquartile range are indicated by circles. Letters groupings indicate communities exhibiting statistically distinct ethylene production (p < 0.05) as determined by the pairwise Games-Howell test following the Levene test for homoscedasticity and the Welch ANOVA (n=10-12). Full statistics for all plots are provided in the Extended Statistics.

To select a community as a starting point for developing a microbial inoculant, we analyzed communities that contained Kv and Hs and lacked Bp in our experimental dataset (**Figure S9**). Higher richness microbial inoculants may be more complex to manufacture. Therefore, we selected a five-member community with relative nitrogenase activity similar to other higher richness communities (**Figure S9)**. This community comprises three diazotrophs (Kv, Hs, and Ss) and two non-diazotrophs (Pp and Ci). We performed an additional experiment with different combinations of this fivd species, examining nitrogenase activity normalized to the total abundance of diazotrophic species (**Figure 5B)**. The subcommunity containing the three diazotrophs had significantly higher nitrogenase activity than each diazotroph in monoculture (**Figure 5B**). Notably, pairs of diazotrophs did not have significantly higher relative nitrogenase activity than monocultures (**Figure S7**), suggesting the presence of higher-order interactions benefiting nitrogenase activity in the three diazotroph community. The presence of Pp or Ci did not significantly influence nitrogenase activity. While in this specific community, nitrogenase activity did not benefit from the presence of non-diazotrophs, the presence of Ci and Pp did not significantly reduce nitrogenase activity (**Figure 5B)**. This three-member diazotroph community of Ss, Hs, and Kv, has higher nitrogenase activity than any of these diazotrophs alone, providing a potential starting point for future microbial inoculant development.

## DISCUSSION

It is commonly assumed that the growth and persistence of organisms that contribute to a given function are a major determinant of the community-level function capacity^26,27^. Our results provide evidence for a more complex scenario. While increased diazotroph growth was positively correlated with nitrogenase activity, the relationship was far from linear, with many communities having higher nitrogenase activity than would be predicted by their diazotroph growth alone. The beneficial interactions observed in pairwise communities were not associated with increased diazotroph growth. For example, both the Ab, Bp pair and the Av, Hr pair had higher nitrogenase activity but the same diazotroph abundance when compared to the diazotrophic monocultures, suggesting growth-uncoupled mechanisms underlying the enhancement in nitrogenase activity. The inferred interspecies interactions were specific to a given diazotroph, rather than generalizable across diazotrophs, consistent with other community functions such as butyrate production in human gut communities^34,38^.

Nitrogenase expression and activity are tightly regulated processes sensitive to oxygen, nitrogen, and energy status, with different diazotrophs favoring different conditions^9,17^. A non-diazotroph could generate a local environment more or less favorable for the fixation of a particular diazotroph by consuming oxygen or nitrogen, or by cross-feeding^30^. The nitrogenase enzyme is a highly complex molecular machine, requiring iron and molybdenum as cofactors^9^. Non-diazotrophs may impact a diazotroph’s ability to access these key micronutrients through changes in extracellular pH or the uptake or release of chelating molecules. The beneficial impact of Bp on the nitrogenase activity of Ab is achieved through modifications to the extracellular environment. An avenue of future research could involve identifying key compounds mediating these interactions. By contrast, the detrimental impact of Bp on the nitrogenase activity of Kv was not reproduced by Bp conditioned media, suggesting that the mechanism requires the two species actively growing together. Our community contained wildtype diazotrophic strains that spanned a range of genetic diversity. However, strains with enhanced nitrogenase activity through deregulated nitrogenase expression may respond differently to community-level interactions, as genetic deregulation alters mutant strains’ sensitivity to their environment^17–20^. Future experiments could explore this further, for example determining whether an engineered strain of KV is still detrimentally impacted by Bp. The broad range of species-specific interactions identified in our dataset presents opportunities for investigation of the mechanisms underlying these patterns.

Three diazotrophs emerged as positive contributors to nitrogenase activity across various community contexts: *Klebsiella variicola* (Kv), *Herbasprillum seropedicae* (Hs), and *Azotobacter vinelandii* (Av). The Kv and Hs strains used in this study were isolated from cereal mucilage and are capable of fixing nitrogen under microaerobic conditions, such as those used in this study^20,43^. By contrast, Av is a soil bacterium that prefers aerobic conditions^42^. Despite this, in our microaerobic experiments, Av fixed nitrogen in coculture with non-diazotroph Hr and contributed positively to nitrogenase activity in most communities, underscoring the benefit of interspecies interactions. In addition to fixing nitrogen directly, diazotrophs may also be able to promote the nitrogenase activity of another fixing species by mechanisms similar to those discussed for non-diazotrophs. For example, it is possible that in some community combinations Av does not need to fix nitrogen directly to be able to positively contribute to the overall nitrogenase activity of the community. Future experiments could use non-fixing mutants to decode these contributions^9,18^.

Previous work developing inoculants for nitrogen-fixation has focused on optimizing or isolating individual diazotrophic strains with high nitrogenase activity that are genetically tractable^21–25^. Our results suggest that strategies focused on two to three diazotrophs are likely to improve nitrogenase activity as opposed to optimizing communities containing a single diazotroph. From a technical perspective, minimizing the number of species in an inoculant is likely to be beneficial. We identified the three-member community containing Ss, Hs, and Kv as a potential useful inoculant because it displayed nitrogenase activity similar to the nitrogenase activity of four- and five-member communities. While in this community we did not observe a benefit to including non-diazotrophs, the presence of these strains did not decrease nitrogenase activity. The positive SHAP contribution of Av suggests that it may also enhance nitrogenase activity in this context. All of these diazotrophs are candidates for microbial inoculants on crops^20,54,55^. Recently, researchers have engineered the nitrogenase activity of several *Klebsiella variicola* strains, including strain A3 used in this study^20^, and successfully applied both wild-type and engineered strains to cereal crops^24,25,29^. With Kv emerging as a key nitrogen-fixing chassis and a major driver of community nitrogenase activity in our study, genetic engineering and consortia development could be combined to optimize nitrogenase activity.

The rhizosphere is a dynamic environment, where metabolites change in response to plant growth, stress and environmental variation^56,57^. Analysis of our experimental dataset revealed a saturating benefit to adding diazotrophic species to a community. The saturation response may arise due to limiting carbon sources or nitrogenase cofactors. A key unresolved question is how this saturating trend shifts as a function of the number or concentration of resources in the environment. Further exploration of these limitations could inform the development of microbial inoculants and complementary strategies such as nutrient or prebiotic supplementation^58,59^. Interactions between strains in microbial inoculants and the resident microbiome also determine the performance of these technologies^26,27^. Within the rhizosphere species included in our study, the growth and nitrogenase activity of Kv was robust across community contexts, consistent with this species’ ability to occupy diverse ecological niches^60^. Future research could explore this robustness further by amending synthetic communities of interest to natural communities^38^.

A detailed and quantitative understanding of microbial interactions and metabolic mechanisms will aid in the development of high performing and robust microbial inoculants that successfully drive these complex systems towards desirable functions^3–5^. Previous efforts to enhance nitrogen fixation in free-living and associative diazotrophs have focused on disrupting negative feedback loops based on nitrogen status as major control knobs^9,17^. This work expands the view to consider the activity of other species as important effectors in this process. Novel metabolic mechanisms that enhance or suppress nitrogen fixation could be elucidated by expanding the diversity of tested species and strains and developing higher throughput assays for quantifying nitrogen fixation activity and using genome-informed computational modeling^61^. Finally, an exciting future direction for this work is the combination of genetic engineering and consortia design to further optimize nitrogen-fixing microbial inoculants.

## Data and code availability

All data and code used for model fitting and experimental design will be publicly available through GitHub (https://github.com/VenturelliLab/).

## Author contributions

C.M.P., J.M.A., and O.S.V. conceived the study. C.M.P. carried out experiments with assistance from J.H.H., and T.W.R. J.T. designed and performed computational modeling. C.M.P. performed statistical analysis of experimental and simulated data. C.M.P., and O.S.V. wrote the manuscript and J.M.A. and J.T. provided feedback on the manuscript.

## Acknowledgments

This research was supported by the Department of Energy under Grant Number DE-SC0021052 (O.S.V. and J.M.A.) and the Agricultural Microbiomes in Plant Systems and Natural Resources, project award number 2023-67012-40302 (C.M.P.), from the U.S. Department of Agriculture’s National Institute of Food and Agriculture. The funders had no role in the study’s conceptualization, data analysis, decision to publish, or preparation of the manuscript.

