## Supplementary Information for "Decoding the role of microbial interspecies interactions on nitrogen fixation"

Table S1: Strains used in this study.

| Species | Strain | Citation |
|---|---|---|
| *Stenotrophomonas maltophilia* (Sm) | AA1 | Niu *et al.*, 2017 |
| *Brucella pituitosa* (Bp) | AA2 | Niu *et al.*, 2017 |
| *Curtobacterium pusillum* (Cp) | AA3 | Niu *et al.*, 2017 |
| *Enterobacter ludwigii* (El) | AA4 | Niu *et al.*, 2017 |
| *Chryseobacterium indologenes* (Ci) | AA5 | Niu *et al.*, 2017 |
| *Herbaspirillum robiniae* (Hr) | AA6 | Niu *et al.*, 2017 |
| *Pseudomonas putida* (Pp) | AA7 | Niu *et al.*, 2017 |
| *Azospirillum brasilense* (Ab) | FP2 | Pedrosa & Yates, 1984 |
| *Azotobacter vinelandii* (Av) | DJ | Setubal *et al.*, 2009 |
| *Stutzerimonas stutzeri* (Ss) | A1501 | Yan *et al.*, 2008 |
| *Herbaspirillum seropedicae* | A5 | Palmer *et al.*, 2025 |
| *Klebsiella variicola* | A3 | Venkataraman *et al.*, 2023 |

Table S2: Media recipes tested in Figure S1.

| **CMP-ARE** | |
|---|---|
| Magnesium sulfate | 0.83 mM |
| Monopotassium phosphate | 0.735 mM |
| Dipotassium phosphate | 2.3 mM |
| Iron (III) chloride | 4.45 µM |
| Sodium molybdate | 0.6 µM |
| Calcium chloride | 0.95 mM |
| Sucrose | 29.2 mM |
| L-arabinose | 9.05 mM |
| Sodium fumarate | 2.9 mM |
| Sodium pyruvate | 1.5 mM |
| Sodium citrate | 350 µM |
| Glutamate | 155 µM |
| Aspartate | 135 µM |
| Alanine | 130 µM |
| Glycine | 80 µM |

| | |
|---|---|
| pH | 6.8 |
| **Medium A** | |
| Magnesium sulfate | 0.5 mM |
| Monopotassium phosphate | 0.7 mM |
| Dipotassium phosphate | 0.8 mM |
| Ferric EDTA | 0.05 mM |
| Manganese sulfate | 1 µM |
| Copper(II) sulfate | 0.7 µM |
| Zinc sulfate | 0.6 µM |
| Boric acid | 1.6 µM |
| Sodium molybdate | 0.5 µM |
| Calcium chloride | 1 mM |
| Sucrose | 29.2 mM |
| L-arabinose | 9.05 mM |
| Sodium fumarate | 2.9 mM |
| Sodium pyruvate | 1.5 mM |
| Sodium citrate | 350 µM |
| Glutamate | 155 µM |
| Aspartate | 135 µM |
| Alanine | 130 µM |
| Glycine | 80 µM |
| pH | 6.8 |
| **Medium B** | |
| Magnesium sulfate | 0.8 mM |
| Monopotassium phosphate | 4.4 mM |
| Dipotassium phosphate | 1.15 mM |
| Iron(II) sulfate | 72 µM |
| Nitrilotriacetic acid | 293 µM |
| Sodium molybdate | 0.2 µM |
| Calcium chloride | 0.18 mM |
| Sucrose | 29.2 mM |
| L-arabinose | 9.05 mM |
| Sodium fumarate | 2.9 mM |
| Sodium pyruvate | 1.5 mM |
| Sodium citrate | 350 µM |
| Glutamate | 155 µM |
| Aspartate | 135 µM |
| Alanine | 130 µM |
| Glycine | 80 µM |

| | |
|---|---|
| pH | 6.8 |
| **Medium D** | |
| Sodium chloride | 1.7 mM |
| Magnesium sulfate | 0.8 mM |
| Dipotassium phosphate | 2.9 mM |
| Iron(II) sulfate | 72 μM |
| Nitrilotriacetic acid | 293 μM |
| Manganese sulfate | 16 μM |
| Copper(II) sulfate | 0.4 μM |
| Zinc sulfate | 0.8 μM |
| Boric acid | 40 μM |
| Sodium molybdate | 10 μM |
| Calcium chloride | 0.18 mM |
| Sucrose | 29.2 mM |
| L-arabinose | 9.05 mM |
| Sodium fumarate | 2.9 mM |
| Sodium pyruvate | 1.5 mM |
| Sodium citrate | 350 μM |
| Glutamate | 155 μM |
| Aspartate | 135 μM |
| Alanine | 130 μM |
| Glycine | 80 μM |
| pH | 6.8 |

Table S3: Preculturing conditions of each species

| **Species** | **First Preculture Timing** | **First Preculture Medium** | **Second Preculture Timing** | **Second Preculture Inoculation Volume** | **Second Preculture Conditions** |
|---|---|---|---|---|---|
| Sm | 19 hours | TY | 24 hours | 100 μL | microaerobic |
| Bp | 19 hours | TY | 24 hours | 100 μL | microaerobic |
| Cp | 19 hours | TY | 24 hours | 100 μL | microaerobic |
| El | 19 hours | TY | 24 hours | 100 μL | microaerobic |
| Ci | 19 hours | TY | 24 hours | 100 μL | microaerobic |
| Hr | 19 hours | TY | 24 hours | 100 μL | microaerobic |

| | | | | | |
|---|---|---|---|---|---|
| Pp | 19 hours | TY | 24 hours | 100 µL | microaerobic |
| Ab | 24 hours | TY+sucrose | 41 hours | 100 µL | microaerobic |
| Av | 41 hours | TY+sucrose | 24 hours | 100 µL | aerobic |
| Ss | 19 hours | TY | 24 hours | 500 µL | microaerobic |
| Hs | 19 hours | TY | 24 hours | 100 µL | microaerobic |
| Kv | 19 hours | TY | 24 hours | 100 µL | microaerobic |

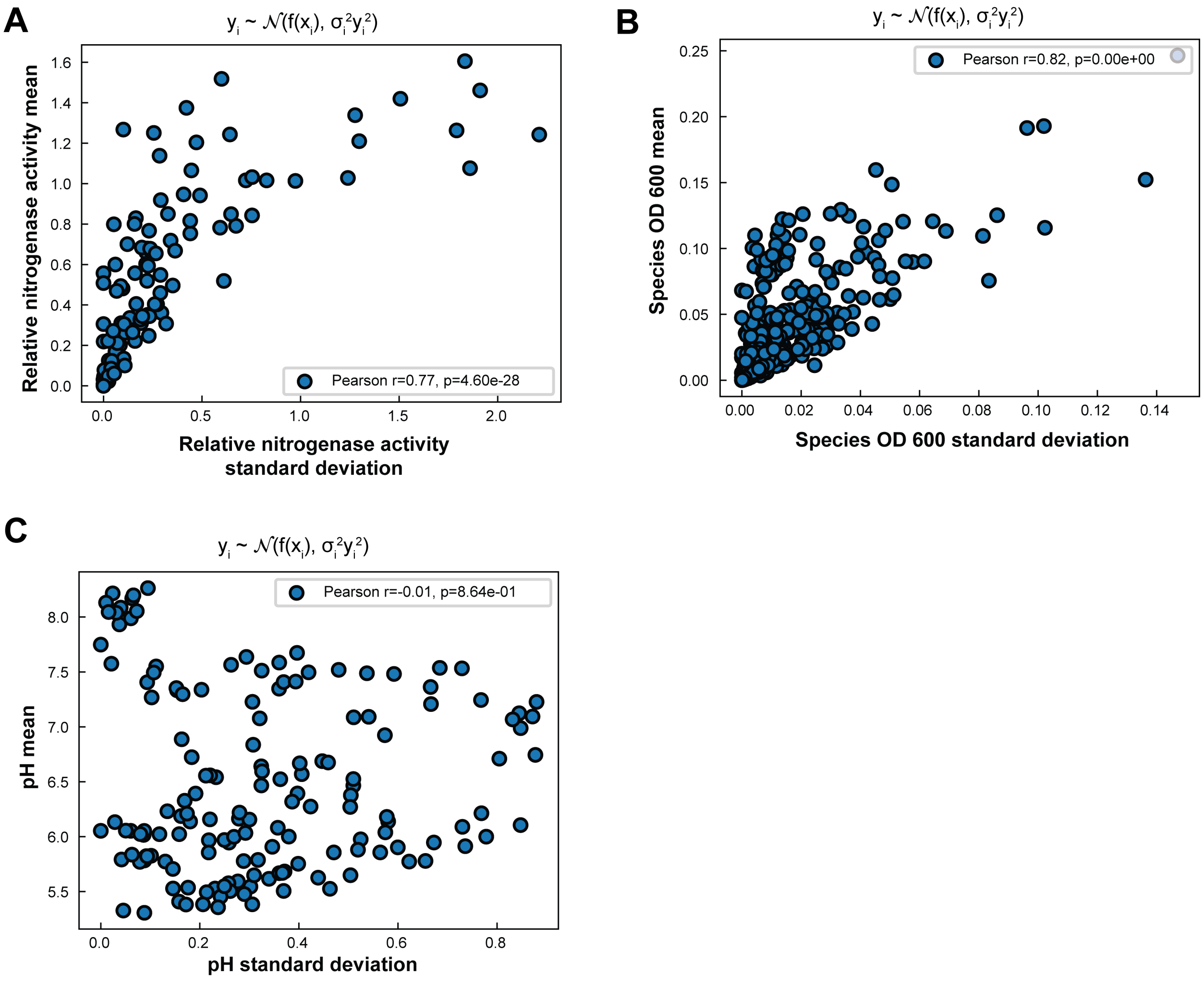


**Figure S1. Relationship between measurement mean and standard deviation across replicates.** A) Measurement noise (standard deviation over measurements of replicates) of relative nitrogenase activity is significantly positively correlated with the mean nitrogenase activity. The correlation between measurement noise and the average value indicates a heteroscedastic process, meaning that the ability to measure a variable depends on its actual value. B) Similar to relative nitrogenase activity, the measurement noise of absolute species abundance (OD 600) is correlated with the average value. C) In contrast, the measurement noise of pH is not correlated with the magnitude of measured pH, consistent with a standard homoscedastic (constant variance) model of measurement variance. Means and standard deviations for all variables were calculated from replicate measurements.

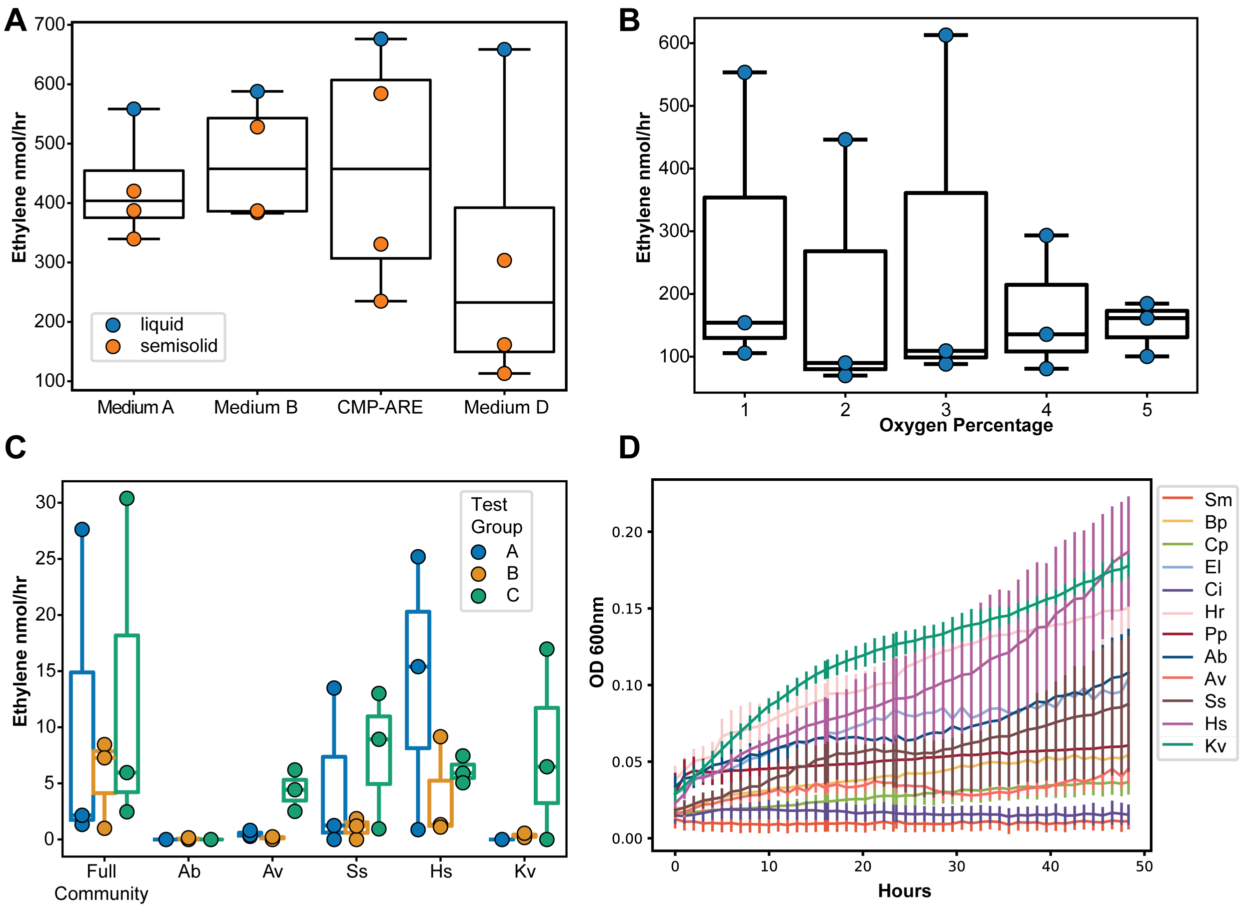


**Figure S2. Selection of experimental conditions.** A) The media outlined in Table S2 were compared by assessing the nitrogenase activity of the full community. Vials containing 3 ml cultures of either semisolid or liquid media were cultured either aerobically (for semisolid media) or in 1% oxygen (for liquid media) for 26.5 hours before adding acetylene. Cultures were then incubated for an additional 17 hours before ethylene concentrations were assessed. Points represent biological replicates. Boxplot hinges indicate the first and third quartiles. B) Different oxygen levels were compared by assessing the nitrogenase activity of the full community in CMP-ARE liquid medium. Vials containing 3 mL cultures were grown for 41 hours before the addition of acetylene. Cultures were incubated for an additional 4 hours before ethylene concentrations were assessed. C) The nitrogenase activity of the full community and each diazotrophic monoculture was assessed in 1 ml cultures of CMP-ARE grown in 3% oxygen. Vials in test group A were cultured for 27 hours before acetylene addition and 17 hours after. Vials in test group B were cultured for 27 hours before acetylene addition and 21.4 hours after. Vials in test group C were cultured for 44 hours before acetylene addition and 4.5 hours after. D) Each species was cultured in monoculture in 200 µL of CMP-ARE. The optical density at 600 nm was recorded hourly. Lines represent the mean of biological replicates, with error bars representing one standard deviation in either direction (n=7).

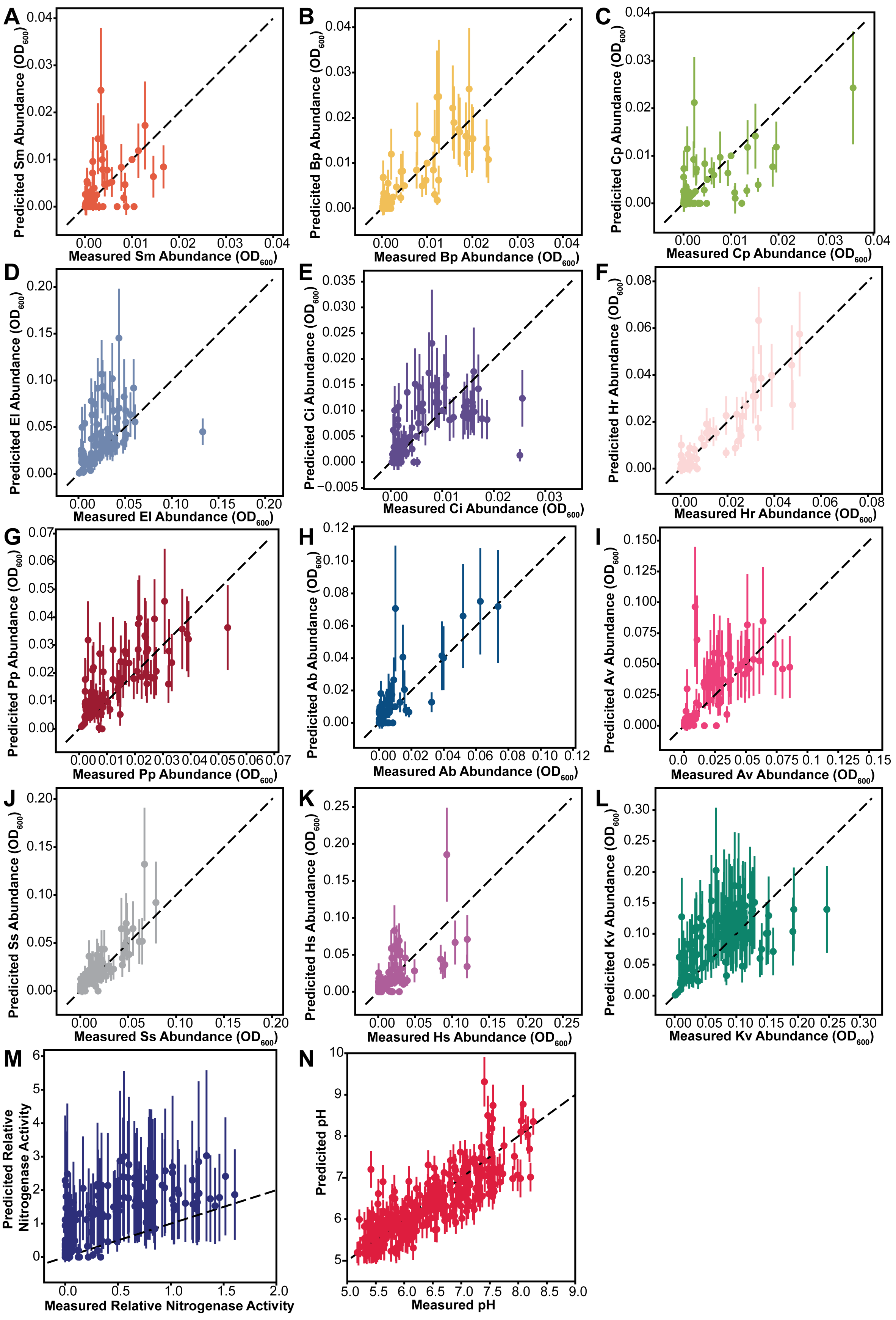
A
Predicited Sm Abundance (OD600)
Measured Sm Abundance (OD600)
B
Predicited Bp Abundance (OD600)
Measured Bp Abundance (OD600)
C
Predicited Cp Abundance (OD600)
Measured Cp Abundance (OD600)
D
Predicited El Abundance (OD600)
Measured El Abundance (OD600)
E
Predicited Ci Abundance (OD600)
Measured Ci Abundance (OD600)
F
Predicited Hr Abundance (OD600)
Measured Hr Abundance (OD600)
G
Predicited Pp Abundance (OD600)
Measured Pp Abundance (OD600)
H
Predicited Ab Abundance (OD600)
Measured Ab Abundance (OD600)
I
Predicited Av Abundance (OD600)
Measured Av Abundance (OD600)
J
Predicited Ss Abundance (OD600)
Measured Ss Abundance (OD600)
K
Predicited Hs Abundance (OD600)
Measured Hs Abundance (OD600)
L
Predicited Kv Abundance (OD600)
Measured Kv Abundance (OD600)
M
Predicited Relative Nitrogenase Activity
Measured Relative Nitrogenase Activity
N
Predicited pH
Measured pH

**Figure S3. Prediction performance of species, relative nitrogenase activity, and pH from 20-fold cross-validation using the MiRNN model.** The entire dataset (averaged across replicates) was split into 20 unique subsets. Each subset was reserved for held-out testing, while the model was trained on the remaining 19 subsets. The training and testing process was repeated 20 times, so that every sample was held out for testing. Each point in the scatter plot shows the model-predicted mean for a test sample, with error bars indicating one standard deviation of the predicted distribution.

Table S4: Prediction performance of the MiRNN model from 20-fold cross-validation. Pearson and Spearman correlation were calculated using Python package scipy. Root mean squared error was calculated using Python package scikit-learn.

| **Variable** | **Pearson r** | **Pearson p-value** | **Spearman r** | **Spearman p-value** | **RMSE** |
|---|---|---|---|---|---|
| Sm Abundance | 0.5677 | 3.4077e-12 | 0.7065 | 1.7205e-20 | 0.0033 |
| Bp Abundance | 0.8364 | 1.3635e-37 | 0.7955 | 1.3179e-31 | 0.0032 |
| Cp Abundance | 0.6964 | 2.1464e-24 | 0.6462 | 3.6135e-20 | 0.0033 |
| El Abundance | 0.6158 | 6.3304e-15 | 0.8448 | 1.4472e-36 | 0.0235 |
| Ci Abundance | 0.6867 | 1.7196e-30 | 0.7198 | 1.1812e-34 | 0.0035 |
| Hr Abundance | 0.9172 | 1.2819e-66 | 0.80678 | 7.4292e-39 | 0.0042 |
| Pp Abundance | 0.7984 | 1.2025e-28 | 0.8146 | 1.1986e-30 | 0.0072 |
| Ab Abundance | 0.8473 | 1.532e-37 | 0.659 | 8.7475e-18 | 0.0075 |
| Av Abundance | 0.8071 | 3.2873e-41 | 0.8214 | 8.575e-44 | 0.0126 |
| Ss Abundance | 0.8933 | 1.5903e-66 | 0.8059 | 3.2954e-44 | 0.0081 |
| Hs Abundance | 0.6869 | 2.9706e-29 | 0.7298 | 1.5562e-34 | 0.0155 |
| Kv Abundance | 0.7965 | 2.2433e-43 | 0.9187 | 1.4284e-78 | 0.0359 |
| pH | 0.848 | 5.1916e-135 | 0.8213 | 1.3815e-119 | 0.3733 |
| Relative Nitrogenase Activity | 0.7912 | 1.8077e-72 | 0.8926 | 3.1715e-116 | 0.695 |

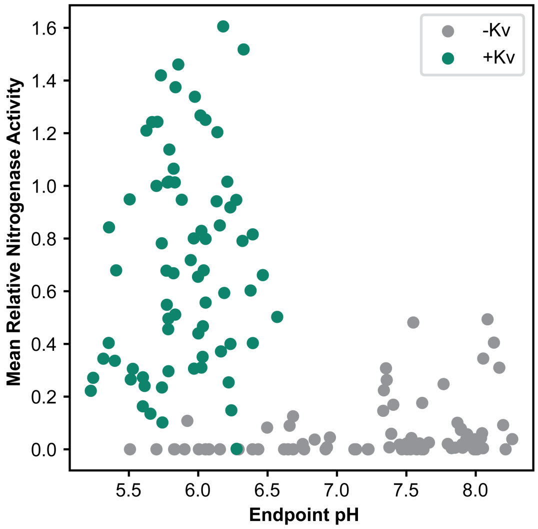


**Figure S4. Relative nitrogenase activity versus pH.** Each datapoint represents a unique community in the experimental dataset (n=149). Color represents the presence/absence of *Klebsiella variicola* (Kv).

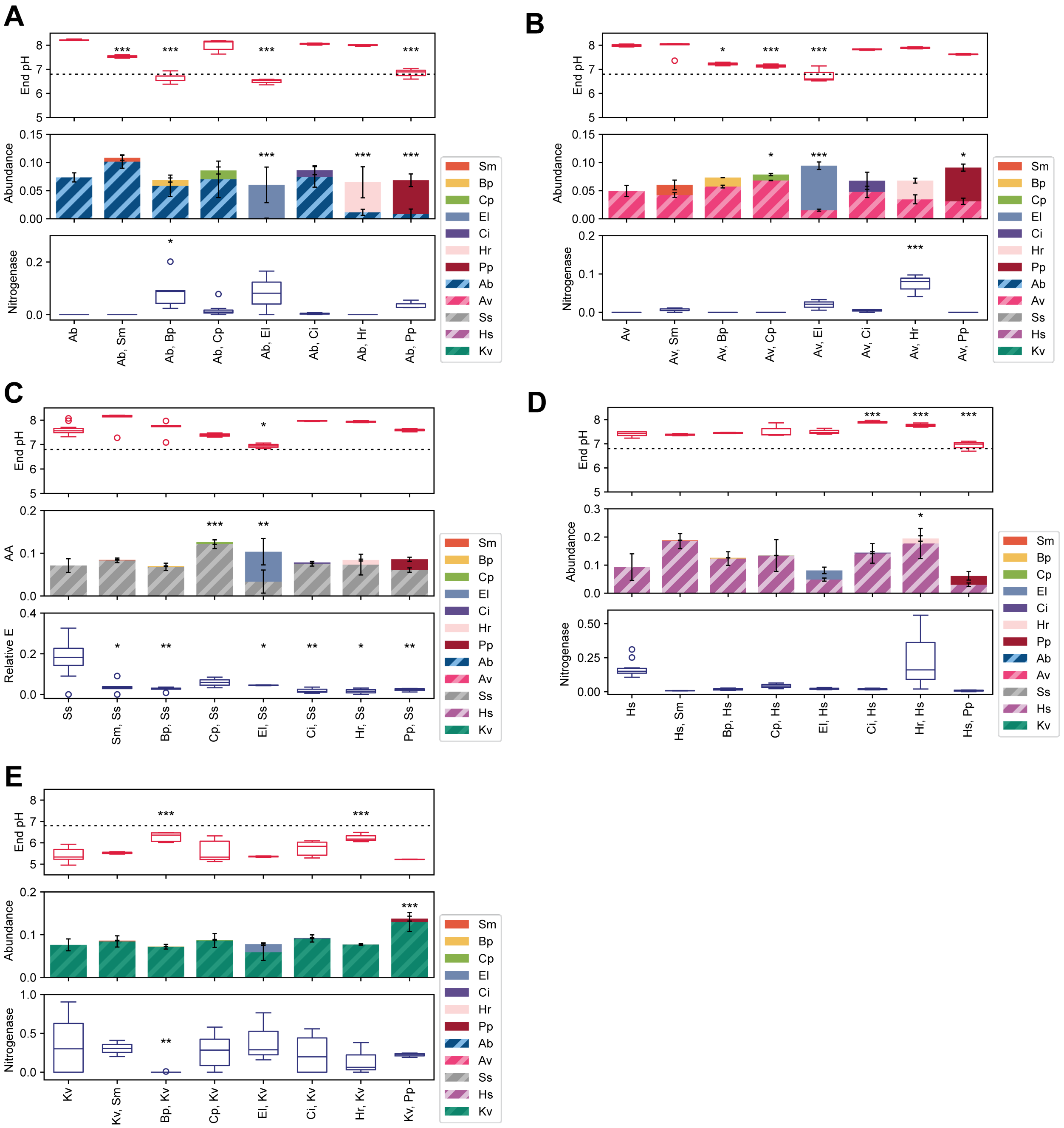

**Figure S5. Impact of non-diazotrophic coculture on each diazotroph.** For each grouping, the top panel shows the endpoint pH determined via phenol red assay, with the dotted line indicating the initial pH of the growth medium. Boxplot hinges indicate the first and third quartiles, and biological replicates outside 1.5 times the interquartile range are indicated by circles. The middle panel represents the community composition after 48 hours, with absolute abundance calculated as the product of 16S relative abundance and cell density at 600 nm. Color represents species, with diazotrophs indicated by hashes. Error bars represent one standard deviation from the mean of biological replicates. The bottom panel shows relative nitrogenase activity. Boxplot hinges indicate the first and third quartiles, and biological replicates outside 1.5 times the interquartile range are indicated by circles. For each diazotroph, pairwise cocultures were compared with the monoculture control using Dunnett's test when variances were equal (determined by the Levene test) and the Games-Howell test when variances were unequal. Full statistics are listed in Extended Statistics. * p < 0.05, ** p < 0.01, *** p < 0.005

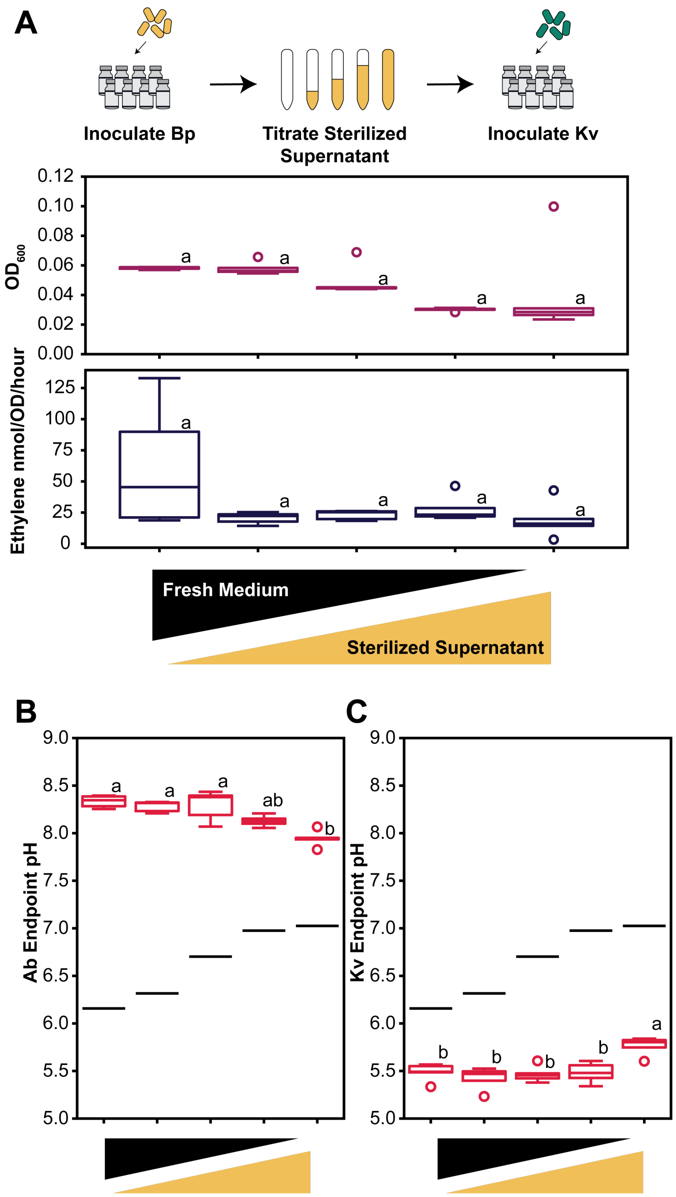


**Figure S6. Impact of Bp sterilized supernatant on diazotrophs Ab and Kv.** A) Growth and ethylene production of Kv monoculture, B) endpoint pH of Ab monoculture and C) endpoint pH of Kv monoculture in (from left to right) 100% fresh medium, 75% fresh medium and 25% Bp sterilized supernatant, 50% fresh medium and 50% Bp sterilized supernatant, 25% fresh medium and 75% Bp sterilized supernatant, and 100% Bp sterilized supernatant. In B) and C) the black lines indicate the pH of a blank of the same medium mixture. Boxplots represent the distribution of biological replicates with hinges indicating the first and third quartiles and circles indicating replicates outside of 1.5 times the interquartile range. Letters represent statistically significant groupings ($p < 0.05$) as determined by Tukey test where a Levene test for equality of variance was true and Games-Howell where it was false (n-5). Full results of all statistical tests are provided in the Extended Statistics table.

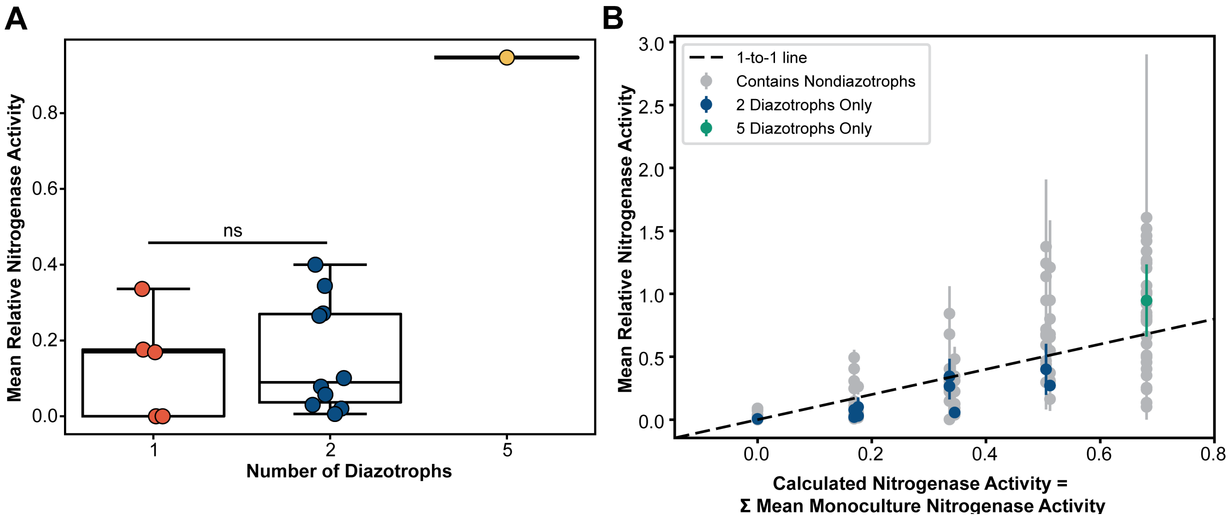


**Figure S7. Impact of multiple diazotroph communities on relative nitrogenase activity.** A) Relative nitrogenase activity of each diazotroph-only community in the experimental dataset grouped by the number of diazotrophic species. Each data point represents a unique community, with color representing the total number of species in the community. Boxplot hinges indicate the first and third quartiles. The letters ns represent a nonsignificant difference ($p < 0.05$) as determined by the pairwise Tukey test following the Levene test for homoscedasticity and ANOVA. Full statistics are listed in Extended Statistics. B) Relative nitrogenase activity of each diazotroph-containing community in the experimental dataset, excluding the monocultures, compared to the sum of monoculture nitrogenase activity of each diazotroph in the community. The dashed line represents a one-to-one relationship. Each data point represents a unique community, with color indicating whether the community contains non-diazotrophic species and, for diazotroph-only communities, the number of species. Error bars represent the standard error of the mean. One standard error below the mean relative nitrogenase activity exceeded the calculated nitrogenase activity in 30.8% of the communities, 149 communities total (n=2-32).

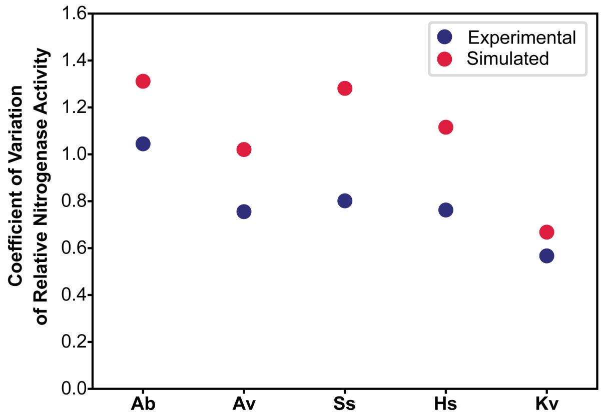


**Figure S8. Diazotroph robustness to community context.** Coefficient of variation of relative nitrogenase activity across communities containing each diazotroph. Each point represents the standard deviation of the mean relative nitrogenase activity for each unique community containing the diazotroph, in either the experimental or simulated datasets, divided by the mean of means.

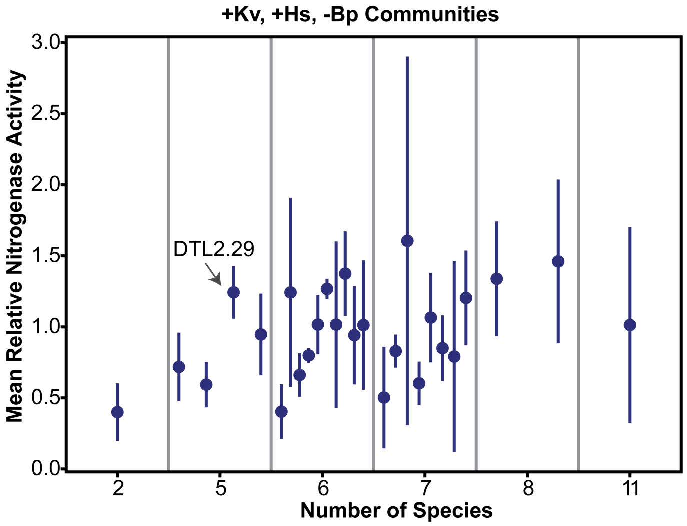


**Figure S9. Selection of a +Kv, +Hs, -Bp community.** The number of species in the community groups communities in the functional dataset containing Kv and Hs and lacking Bp. Each datapoint represents the mean of a unique community, with error bars representing the standard error of the mean. The community of interest is indicated with an arrow (n=2-17).
